# Nucleus- and plastid-targeted annexin 5 promotes reproductive development in Arabidopsis and is essential for pollen and embryo formation

**DOI:** 10.1101/331249

**Authors:** Malgorzata Lichocka, Wojciech Rymaszewski, Karolina Morgiewicz, Izabela Barymow-Filoniuk, Aleksander Chlebowski, Miroslaw Sobczak, Marcus A. Samuel, Elmon Schmelzer, Magdalena Krzymowska, Jacek Hennig

**Affiliations:** Institute of Biochemistry and Biophysics, Polish Academy of Sciences, Pawinskiego 5a, 02-106 Warsaw, Poland; Institute of Biochemistry and Biophysics, Polish Academy 20 of Sciences, Pawinskiego 5a, 02-106 Warsaw, Poland; Department of Botany, Warsaw University of Life Sciences (SGGW), Warsaw, Poland; Department of Biological Sciences, University of Calgary, Alberta, Canada; Max-Planck Institute for Plant Breeding Research, Cologne, Germany

**Keywords:** Arabidopsis, accession, annexin, pollen grain, seed, embryo, plastid, nucleoid, chlorophyll, Rab GTPase

## Abstract

**Background:** Pollen development is a strictly controlled post-meiotic process during which microspores differentiate into microgametophytes and profound structural and functional changes occur in organelles. Annexin 5 is a calcium- and lipid-binding protein that is highly expressed in pollen grains and regulates pollen development and physiology. To gain further insights into the role of ANN5 in Arabidopsis development, we performed detailed phenotypic characterization of Arabidopsis plants with modified *ANN5* levels. In addition, interaction partners and subcellular localization of ANN5 were analyzed to investigate potential functions of ANN5 at cellular level.

**Results:** Here, we report that RNAi-mediated suppression of *ANN5* results in formation of smaller pollen grains, enhanced pollen lethality, and delayed pollen tube growth. *ANN5* RNAi knockdown plants also displayed aberrant development during the transition from the vegetative to generative phase and during embryogenesis, reflected by delayed bolting time and reduced embryo size, respectively. At the subcellular level, ANN5 was delivered to the nucleus, nucleolus, and cytoplasm, and was frequently localized in plastid nucleoids, suggesting a likely role in interorganellar communication. Furthermore, ANN5-YFP co-immunoprecipitated with RABE1b, a putative GTPase, and interaction *in planta* was confirmed in plastidial nucleoids using FLIM-FRET analysis.

**Conclusions:** Our findings let us to propose that ANN5 influences basal cell homeostasis via modulation of plastid activity during pollen maturation. We hypothesize that the role of ANN5 is to orchestrate the plastidial and nuclear genome activities via protein-protein interactions however not only in maturing pollen but also during the transition from the vegetative to the generative growth and embryo development.

## Background

In angiosperms, the male gametophyte (microgametophyte or pollen grain) plays an essential role in the reproductive success of the species, and normal pollen development under challenging environmental conditions is a highly desirable agronomic trait in various crops. Development of a male gametophyte is a complex process that takes place in the anther locules, where microspore mother cells undergo meiosis to produce haploid microspores [1, 2]. Developing microspores take up nutrients from the tapetum, an inner layer of cells in the anther locule. This secretory tissue provides soluble carbohydrates for microspore growth and lipids for pollen cell wall formation [3, 4]. Despite being dependent upon nutrient delivery from the tapetum, microspore plastids undergo intensive structural reorganization as the microspore matures [2]. In young microspores, plastids are poorly differentiated and lack any internal membranous system. Before the first mitosis, the plastids develop a few thylakoids and differentiate into amyloplasts and accumulate starch transiently until the bicellular stage of microgametogenesis. Following the second mitosis, the tricellular mature pollen grain is made up of one vegetative cell (VC) and two sperm cells. At this stage, the pollen plastids contain only negligible amounts of starch as the majority of the starch is hydrolyzed [5]. Although limited in number in developing microspores, these plastids are crucial for pollen viability as various mutants defective in plastid carbohydrate metabolism exhibit pollen sterility [6].

Genes important for male gametophyte development can be assigned as either ‘early’ or ‘late’, according to their spatiotemporal expression pattern. The ‘early’ genes are the first to be activated in the microspore, and their expression levels decrease as pollen maturation approaches. The ‘late’ genes are activated after the first microspore mitosis, and their transcripts accumulate during pollen maturation [7].One of the late genes in the developing microspore is annexin 5 (*ANN5*). *ANN5* promoter activity was detected in the bicellular microspore, and maximum *ANN5* transcript abundance correlated with pollen maturation [7, 8]. Annexins belong to a ubiquitous family of proteins present in eukaryotic organisms [9, 10] localized to various subcellular compartments [11]. Due to their calcium- and membrane-binding capacity, annexins are known to be involved in a variety of cellular processes such as actin binding, maintenance of vesicular trafficking, cellular redox homeostasis, and ion transport [12]. ANN5 was previously characterized biochemically and, like other annexins, associated with liposomes in a calcium-dependent manner and bound actin [13]. Pollen tubes overexpressing *ANN5* displayed enhanced resistance to Brefeldin A (BFA), an inhibitor of vesicular protein transport, which suggested that ANN5 promoted membrane trafficking downstream of the block by BFA. Supporting this, RNAi-based down-regulation of *ANN5* resulted in enhanced pollen lethality [8]. However, the mechanisms through which ANN5 affects microspore development remain unknown. Our results show that ANN5 function is not limited to male gametophyte development but plays a central role during the entire reproductive development process in Arabidopsis. We further show that ANN5 localizes to the nucleus and the plastids, implicating ANN5 in crosstalk between cellular compartments essential for the maintenance of cellular homeostasis.

## Materials

### Plant material and growth conditions

The experiments were carried out on *Arabidopsis thaliana* and *Nicotiana benthamiana* plants. Modified *ANN5* expression was introduced in Arabidopsis Col-0 background. The other Arabidopsis accessions: An-1, Bay-0, C24, Ler-1, Mr-0, Oy-0 and Wa-1 were obtained from NASC (http://arabidopsis.info/). Arabidopsis plants were grown in Jiffy7 pots in controlled-environment chambers (Percival Scientific, Iowa, USA) at 22°C under 8 h of light and 40% humidity. *N. benthamiana* plants were grown in soil under controlled environmental conditions (21^°^C, 16 h of light).

### Arabidopsis phenotype characterization

For phenotypic studies we used seeds of two selected *ANN5*-RNAi lines: *ANN5*-RNAi_13, *ANN5*-RNAi_15, OE_2 line and wild-type Arabidopsis Col-0. Five seeds of each genotype were placed per Jiffy7 pot. After 2 days of stratification at 4°C the pots were placed under 12 h light/12 h dark photoperiod or under short day (8 h of light) for 4 weeks followed by long day (16 h of light) conditions (sd/ld) at 22°C and 40% humidity in the growth chamber. The light during the day period was provided with mixed fluorescent tubes and incandescent bulbs. Total photon flux density at the soil level was 120 μE m^-2^ s^-1^. After reaching two cotyledons stage only one seedling per pot was further cultured and the rest was removed. Each developmental stage was recorded for 7-10 individual plants per genotype. All plants were daily inspected from germination until siliques ripening. Bolting time was measured as the number of days from germination to the first elongation of the floral stem at 0.1 cm height. Flowering time was estimated as the number of days from germination to the first flower opening. After fading of the first flower the time of silique formation was recorded. Trays with growing plants were rotated three times per week for uniform plant development. Data were analyzed using Microsoft Excel and R freeware software (http://www.r-project.org).

### Seed size measurements

During Arabidopsis growth all the auxiliary buds were removed. Once the siliques formed on the main bolt, turned almost completely brown they were harvested into a microcentrifuge tubes. The siliques were let to air-dry in the open tubes for several days prior to measurements. The dry seeds were dispersed on microscope slides and several images were collected under the stereoscopic microscope (SMZ1500, Nikon Instruments B.V. Europe, Amsterdam, The Netherlands). Following the conversion of images to black-white images using the threshold function of ImageJ software the area of the individual seed was calculated as described previously [14].

### Chlorophyll extraction and measurement

Col-0 wild-type, *ANN5*-OE_2 and *ANN5*_OE1 seeds were surface-sterilized with 75% ethanol for 2 min and then with 10% sodium hypochlorite for 10 min. Next, the seeds were washed three times with sterile water and spread onto agar-solidified (1% w/v) MS media (Duchefa, Amsterdam, The Netherlands) supplemented or not with 1.5% (w/v) sucrose. After 2 days stratification at 4°C, the plates were placed in the growth chamber under a photon flux density of 220 μE m^-2^ s^-1^ at the shelf level. Seedlings were grown under short-day conditions (8 h light) for 10 days. Aerial part of 10 day old seedlings were harvested, weighed and kept at -80°C. Samples were mechanically ground in 2 ml microfuge tubes with two stainless-steel beads by a bead mill (TissueLyser II, Qiagen, Hilden, Germany). After extraction with 1 ml cold 80% acetone, the samples were centrifuged 6000 rpm for 5 min at 4°C. Extraction was repeated two times with fresh solvent. Absorbance of the pooled extracts was measured at 664 and 647 nm with a spectrophotometer (UV-1202, Shimadzu, Kyoto, Japan). Chlorophyll content was calculated using equations described previously [15].

### Plasmid constructions for transient and stable expression in planta

Coding sequence of *ANN5* was PCR amplified using primers adding BglII-BamHI restriction sites: forward 5’-AGATCTCGATGGCGACTCTTAAGGTTTCT-3’ and reverse 5’-GGATCCTAGCATCATCTTCACCGAGAA-3’ and cloned into modified pSAT4A plasmid bearing the full-length cDNA sequence of YFP. The expression cassette 35S:ANN5-YFP was subcloned into pPZP-RCS2 binary plasmid [16]. In parallel, coding sequence of *ANN5* was also PCR amplified with primers adding SalI-EcoRV restriction sites: forward 5’-GTCGACATGGCAACAATGAA-3’ and reverse 5’-GATATCCAACGTTGGGGCCTAAAAGAGAGAG-3’ and cloned into pENTR1A vector compatible with the Gateway system. The resulting plasmid was LR recombined into GWB441 and GWB442 binary plasmids [17]. Coding sequence of *RABE1b* was PCR amplified using primers adding SalI-XhoI restriction sites: forward 5’-GTCGACATGGCGAAGATGATGATGTTGC-3’and reverse CTCGAGGCTTGAAGAACAAGTTTCTTGCTCAG-3’. The amplified coding sequences were cloned into pENTR1A vector, then LR recombined into GWB444 [17].

Agrikola binary plasmids (http://www.agrikola.org/) for targeted *ANN5* RNAi silencing were obtained from NASC (http://arabidopsis.info/) [18]. pAgrikola plasmids contain a fragment of a gene coding sequence, called gene specific tag (GST), under the control of 35S promoter that enables production of double-stranded hairpin RNA (hpRNA) necessary for targeted gene silencing [18]. GST in pAgrikola 35S:*ANN5*(GST)-RNAi corresponded to 214 bp long fragment of *ANN5* coding sequence starting at position 668 and ending at 881.

Wild-type Arabidopsis Col-0 plants were transformed with the following constructs: pPZP-RCS2 *35S:ANN5-YFP*, pAgrikola *35S:ANN5*(GST)-RNAi and pCAMBIA 1302 35S:*GFP* using floral dipping method [19] with *Agrobacterium tumefaciens* strain GV3101 carrying helper plasmids pMP90 and pSOUP. *ANN5*-RNAi transformants were identified using Basta-based selection procedure (http://www.agrikola.org/), whereas selection of 35S:*ANN5-YFP* and 35S:*GFP* transformants was performed directly on MS plates under fluorescence stereo microscope.

To study subcellular localization of ANN5 and RABE1b *Agrobacterium* cultures carrying appropriate constructs were infiltrated into leaves of *N. benthamiana,* which were examined after 72 h under fluorescence confocal microscope (Nikon C1 Instruments B.V. Europe, Amsterdam, The Netherlands).

### RNA extraction and RT-qPCR

Total RNA was isolated from vegetative and generative Arabidopsis tissues using Syngen Plant RNA Mini Kit (Syngen, Wroclaw, Poland). Mature pollen grains were collected on ice-cold 0.3 M mannitol, according to the procedure described previously [20]. Isolated RNA was quantified with a NanoDrop ND-1000 spectrophotometer (Thermo Fisher Scientific, USA) and subjected to DNA digestion (Rapid out DNA removal kit, Thermo Fisher Scientific). First cDNA was synthesized using 2 μg RNA and Superscript III kit (Thermo Fisher Scientific, USA). qPCR was performed with the SYBR green master mix (Thermo Fisher Scientific, USA) using Light Cycler 480 (Roche, Basel, Switzerland). Reactions were run in triplicate with three different cDNA preparations. The relative expression level was normalized with the expression of the reference genes (*UBC21, PP2A* and *YLS8*) and quantified by ΔCt method. Primers for RT-qPCR are listed in Additional file 9.

### Protein extraction and immunoprecipitation

The samples collected from 12 days old Arabidopsis seedlings revealing constitutive expression of ANN5-YFP or GFP were ground in liquid nitrogen. Samples were then thawed in 2 ml of extraction buffer (50 mM Tris-HCl, pH 7.5; 50 mM NaCl; 6 mM EDTA; protease inhibitor PMSF; 0.5% [v/v] Triton X-100) per 1 g of tissue powder. Samples were centrifuged at 13 000 rpm and 4°C for 20 min. Collected supernatants were adjusted to 3 mg ml-1 of total proteins and incubated with GFP-TrapA-beads (Chromotek, USA) for 4 h at 4°C. After incubation the supernatant was discarded and the beads were washed using 50 mM Tris-HCl (pH 7.5), 150 mM NaCl and 2 mM EDTA buffer. Proteins were eluted using 200 mM glycine (pH 2.5). The eluted proteins were trypsin digested and subjected to mass spectrometry.

### Mass Spectrometry Analysis

Liquid chromatography-mass spectrometry analyses of the peptide mixtures were performed on the Orbitrap spectrometer (Thermo Fisher Scientific, USA) and the Mascot program was used for database searches as described previously [21].

### Confocal laser scanning microscopy

Subcellular localization of the fusion proteins was evaluated using a Nikon C1 confocal system built on TE2000E and equipped with a 60× Plan-Apochromat oil immersion objective (Nikon Instruments B.V. Europe, Amsterdam, The Netherlands). GFP/YFP fusion proteins were excited with a Sapphire 488 nm laser (Coherent, Santa Clara, CA, USA) and observed using the 515/530 nm emission filter. CFP fusion protein and DAPI fluorescence were excited with a 408 nm diode laser and detected using the 450/35 nm emission filter. Embryos stained with FM4-64 were excited with 543 nm HeNe laser and observed using the 605/75 nm barrier filter. Confocal images were deconvoluted and pseudocolored using ImageJ software.

### FLIM-FRET

For FLIM-FRET (Fluorescence Lifetime Imaging Microscopy-Förster Resonance Energy Transfer) RABE1b was fused to CFP (donor) and transiently expressed in *N. benthamiana* leaves in the presence or absence of the potential interacting partner ANN5 fused to YFP (acceptor). Cells were imaged with an FV100 confocal system (Olympus, Tokyo, Japan) equipped with a 60x water immersion objective lens. For FLIM CFP fusion protein was excited with a 440 nm pulsed diode laser (Sepia II, PicoQuant, Berlin, Germany) and detected using a 482/35 bandpass filter. Images were acquired with a frame size of 256 × 256 pixels. Photons were collected with a SPAD detector and counted with the PicoHarp 300 TCSPC module (Picoquant). The obtained data were analyzed with Symphotime software (PicoQuant). Fluorescence lifetimes of CFP in plastid nucleoids were calculated by fitting a bi-exponential decay model.

### Transmission and scanning electron microscopy

Flower buds and flowers at anthesis were sampled from Arabidopsis Col-0 wild type, *ANN5*-RNAi_13, *ANN5*-RNAi_15 and *ANN5*-OE_2 genotypes. Plant samples were fixed in a mixture of 2% paraformaldehyd (w/v) and 2% glutaraldehyde (v/v) in 0.05 M sodium cacodylate buffer for 2 h at room temperature. Samples were post fixed in osmium tetroxide, dehydrated in ethanol and embedded in EPON resin according to.[22]. Ultrathin sections were examined in an FEI 268 D ‘Morgagni” (FEI Corp., Hillsboro, OR, USA) transmission electron microscope equipped with an SIS ‘Morada’ digital camera (Olympus SIS, Münster, Germany).

Mature pollen grains were collected directly into a cap of the microfuge tube. They were processed for scanning electron microscopy as described previously [23]. Imaging was performed with a Zeiss Spura 40VP (Zeiss, Jena, Germany) scanning electron microscope operating at 10 kV.

## Results

### *ANN5* is expressed in a tissue-specific manner

Earlier studies demonstrated that *ANN5* was predominantly expressed in mature flowers [8, 24]. To further characterize *ANN5* expression, we analyzed vegetative and reproductive organs of Arabidopsis using RT-qPCR. *ANN5* transcripts were less abundant during vegetative growth than in reproductive tissues of Arabidopsis (Fig. 1A), and were nearly undetectable in 3-day-old seedlings, rosette leaves, and roots. After the transition to the generative phase, a slight increase in *ANN5* expression level was observed in the developing stem and strong expression was detected in young developing siliques. A separate analysis of the pistils and stamens revealed that the strongest *ANN5* expression was seen in the male organs, with the highest *ANN5* transcript abundance observed in mature pollen (Fig. 1B).

**Fig. 1.**
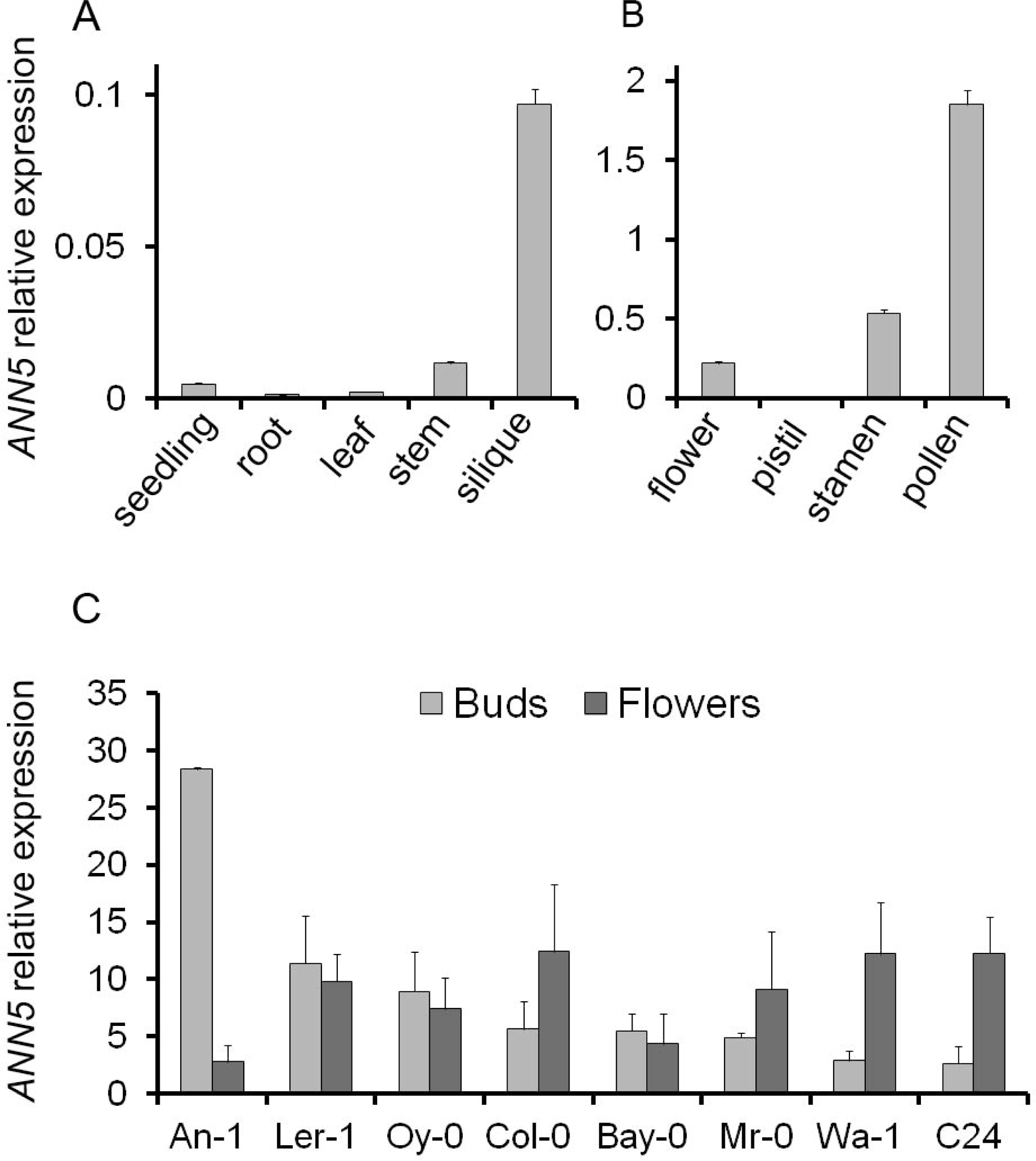
*ANN5* expression profiles. (A)Average expression of *ANN5* in different organs of wild-type Arabidopsis Col-0. (B)Average expression of *ANN5* in reproductive structures of wild-type Arabidopsis Col-0. (C)Average expression of *ANN5* in floral buds and flowers at anthesis collected from different Arabidopsis genotypes. n = 3 biological replicates. Bars represent SD.

### Arabidopsis accessions differ in *ANN5* expression

Elevated expression of *ANN5* correlated with pollen grain maturation in Col-0 plants (Fig. 1B), and we wished to determine whether this was consistent among Arabidopsis accessions. Eight Arabidopsis accessions, originally derived from different habitats, were selected and cultivated until flowering under short day and long day conditions: An-1, C24, Col-0, Ler-1, Bay-0, Wa-1, Oy-0, Mr-0. RT-qPCR analysis of RNA isolated from flower buds and mature flowers revealed differences in *ANN5* expression among the accessions (Fig. 1C). Wa-1, C24, and Mr-0 exhibited a Col-0-type expression pattern with higher *ANN5* mRNA levels in mature flowers than in buds. In Bay-0, Oy-0, and Ler-1, *ANN5* expression was already elevated in the flower buds and remained at similar levels during anthesis. An-1 exhibited the most unusual *ANN5* expression profile: very high expression was observed in flower buds but expression dropped precipitously in the mature flowers. Accession-specific patterns of *ANN5* expression might reflect possible differences in male gametophyte development among Arabidopsis accessions.

### Suppression of *ANN5* leads to a delay in generative development in Arabidopsis

Previous research showed that RNAi (RNA interference)-based suppression of *ANN5* driven by the pollen-specific promoter *LAT52* led to enhanced pollen lethality [8], suggesting that a knockout might be lethal or male sterile. Here, an RNAi approach was used to generate *ANN5* knockdowns using the AGRIKOLA RNAi plasmid carrying 214 bp of the *ANN5* coding sequence under the control of a 35S promoter [18]. The obtained RNAi lines exhibited moderate suppression at anthesis, with *ANN5* levels reduced by 20–80% compared with control Col-0 plants. Two *ANN5* RNAi lines, *ANN5*-RNAi_13 and *ANN5*-RNAi_15, with 50% and 70% *ANN5* suppression, respectively, were selected for detailed phenotypic analysis (Additional file 1). Lines were also generated that ectopically overexpressed *ANN5* (OE) under the control of the 35S promoter. OE lines exhibited extremely elevated *ANN5* transcript abundance compared with the wild type (approximately 100-fold increase) (Additional file 1).

Developmental and morphometric analyses of the selected *ANN5* RNAi lines were conducted under two light regimes: i) 28 days under short day conditions (sd; 8 h light) followed by growth under long day conditions (ld; 12 h light) for the remainder of the experimental period, and ii) 12 h light / 12 h dark throughout the whole experimental period. Germination of all selected lines was equivalent under both conditions. Developmental differences between the RNAi lines and wild-type Col-0 plants became apparent during the transition from vegetative to generative development, i.e., at bolting (formation of a flower stem) (Table 1, Additional file 2). *ANN5* RNAi-silenced lines bolted approximately 8 days later than wild-type plants under the sd/ld light regime (Table 1). By contrast, the bolting delay was only approximately 2 days under the 12 h light regime (Additional file 2). Delay in progression to subsequent growth stages was observed in the *ANN5* RNAi lines, but only under sd/ld conditions. Initiation of flowering and first silique formation were delayed by approximately 11 days in *ANN5* RNAi plants compared with control plants. Transgenic Arabidopsis overexpressing *ANN5* exhibited slightly increased growth rates during rosette formation and stem elongation compared with control plants (Additional file 2); however, developmental progression to the subsequent growth stages was similar to that of wild-type plants under both light regimes (Table 1, Additional file 2).

**Table 1.** Timing of reproductive development in Arabidopsis genotypes with altered *ANN5* expression.

| Genotypes | Bolting |  |  | Flowering |  |  | Silique formation |  |  |
| --- | --- | --- | --- | --- | --- | --- | --- | --- | --- |
|  | [days] |  |  |  |  |  |  |  |  |
| Col-0 | 28.58 | ± | 0.92 | 34.67 | ± | 1.26 | 38.08 | ± | 1.17 |
| ANN5-RNAi_13 | 35.67<sup>a</sup> | ± | 0.89 | 44.83<sup>a</sup> | ± | 1.36 | 49.17<sup>a</sup> | ± | 1.27 |
| ANN5-RNAi_15 | 36.08<sup>a</sup> | ± | 0.62 | 47.10<sup>a</sup> | ± | 0.63 | 50.25<sup>a</sup> | ± | 0.84 |
| ANN5-OE_2 | 30.09 | ± | 1.91 | 35.45 | ± | 1.53 | 38.82 | ± | 1.44 |

Plants were cultivated under a short day/long day regime. Values represent days after germination ± standard error, n = 7 individual plants per line. <sup>a</sup> denotes statistically significant difference (p < 0.05 Dunnett test). See also Additional file 2.

### Pollen viability and grain size correlate with *ANN5* expression level

Phenotypic studies revealed that onset of the generative stage was delayed in *ANN5* RNAi knockdown plants compared with wild-type Col-0 plants, but plant morphological characteristics, i.e., foliage rosette formation, leaf morphology, and inflorescence structure, were generally unaffected. However, abnormal flowers with additional petals and/or missing stamens were observed in *ANN5* RNAi-silenced plants (Additional file 3). We next tested whether suppression of *ANN5* expression affected pollen viability, using Alexander’s solution to differentiate between aborted and non-aborted pollen grains. Anthers of *ANN5* RNAi-silenced lines contained numerous aborted pollen grains (green-colored) and fewer vivid pollen grains (pink-colored) than wild-type and *ANN5*-OE_2 plants (Additional file 3).

Pollen grains from *ANN5* RNAi-silenced lines were examined further using scanning electron microscopy. Cell wall formation was unaffected in *ANN5* RNAi-silenced and *ANN5*-OE_2 pollen, but mean pollen grain size was affected. *ANN5* RNAi-silenced pollen grains were significantly shorter (average 26 μm along the longer axis) than *ANN5*-OE_2 pollen grains (29 μm) and wild-type pollen grains (27.5 μm) (Fig. 2a and Fig. 2C). *ANN5* transcript abundance was previously shown to correlate with pollen maturation in Col-0 plants (Fig. 1C) [7], and we therefore examined pollen maturation in altered and wild-type lines using transmission electron microscopy. Micrographs of pollen grains collected just before and during anthesis showed that progression of pollen grain maturation was similar in all the Arabidopsis genotypes examined (Fig. 2D, Additional file 4). Profound reorganization of the VC encompassed i) partial hydrolysis of starch grains deposited within plastids, ii) formation of numerous initially small vesicles that eventually produced elaborate structures forming ‘foamy’ cytoplasm, and iii) conversion of storage lipids deposited in oil bodies. *ANN5*-RNAi pollen grains only occasionally displayed reduced VC cytoplasm vesiculation compared with control Col-0 pollen. In contrast with wild-type pollen grains, which usually contained a single starch grain per plastid, plastids of *ANN5* RNAi-silenced pollen grains often contained several starch grains. Collapsing pollen grains of *ANN5* RNAi knockdown plants (particularly those of the *ANN5*-RNAi_15 line) contained starch grains that were significantly larger and more numerous than those in aborted pollen of wild-type and *ANN5*-OE_2 plants (Fig. 2E). The high starch content in the collapsing pollen grains of *ANN5* RNAi-silenced lines indicated that abortion of the microspores likely occurred before starch hydrolysis, which normally takes place at the bicellular stage of microgametophyte development.

**Fig. 2.**
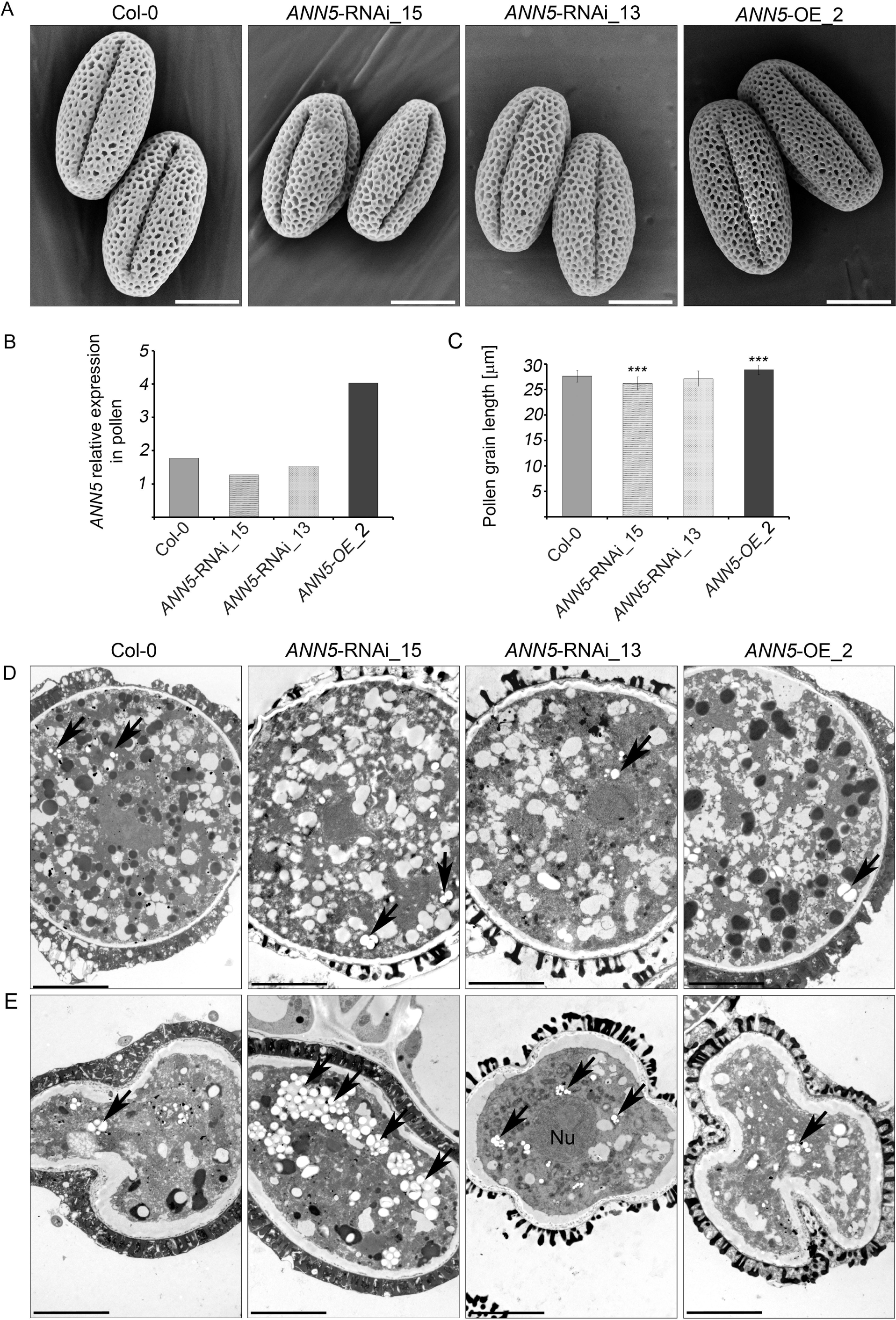
Impact of *ANN5* expression on mature pollen grain size. (A)Scanning electron micrographs of the pollen grains from wild-type Arabidopsis (Col-0), *ANN5*-RNAi_15, *ANN5*-RNAi_13, and *ANN5*-OE_2. Scale bars = 10 μm. (B) Relative expression of *ANN5* in mature pollen grains of wild-type Arabidopsis (Col-0), *ANN5*-RNAi_15, *ANN5*-RNAi_13, and *ANN5*-OE_2. Bars represent SD. (C) Mean length of mature pollen grains from wild-type Arabidopsis (Col-0), *ANN5*-RNAi_15, *ANN5*-RNAi_13, and *ANN5*-OE_2. n = 50. Asterisks indicate significant difference compared with values for wild-type pollen (one-way ANOVA, Dunnett post hoc test, *p < 0.05; **p < 0.01; ***p < 0.001). Bars represent SD. (D)and (E) Ultrastructure of viable and collapsing pollen grains from Arabidopsis genotypes with altered *ANN5* expression. (D) Transmission electron micrographs showing ultrastructural details of viable mature pollen grains, whereas (E) depicts aborted pollen grains isolated during anthesis from wild-type Arabidopsis Col-0, *ANN5*-RNAi_15, *ANN5*-RNAi_13, and *ANN5*-OE_2. See also Additional file 3 and Additional file 4. Nu: nucleus, black arrow: plastid. Scale bars = 5 μm.

### *ANN5* is required for pollen tube growth in pistils

Pollen grain size and viability were affected by altered expression of *ANN5*. Previous research showed that germination rates and pollen tube growth of *ANN5* RNAi-silenced, OE, and wild-type pollen on a solid medium were similar and that the tubes were free of morphological aberrations [13]. Here, hand-pollination of pistils was used to assess the ability of *ANN5* RNAi-silenced and OE pollen grains to germinate and elongate under natural conditions on stigmas. The pistils were collected 3, 6, and *24* hours after pollination and examined for pollen tube growth using a fluorescent technique. Great variations in the growth rate were repeatedly observed among individual pollen tubes derived from the pollen grains of the same *ANN5* RNAi-silenced line. In contrast, the growth rate of pollen tubes in wild-type and OE line were more equivalent. At 3 and 6 h after pollination the majority of the *ANN5* RNAi-silenced pollen tubes did not enter the pistil transmitting tissue although an excess of pollen grains was applied. At 6 h after pollination, pollen from *ANN5* RNAi knockdown plants exhibited shorter pollen tubes (average 0.67 mm from the top of style to the front of the longest pollen tube) than pollen from wild-type (0.88 mm) and *ANN5*-OE_2 line (0.91 mm) (Fig. 3). However, this discrepancy was no longer observed after 24 h (Additional file 5), by which time pollen tubes in all genotypes had traversed to the ovary and reached the ovules. Pollen tube growth rate is a major determinant of pollen competitive ability, and the arrested or delayed growth of *ANN5* RNAi-silenced pollen tubes in the pistil is indicative of lower male gametophyte competitiveness.

**Fig. 3.**
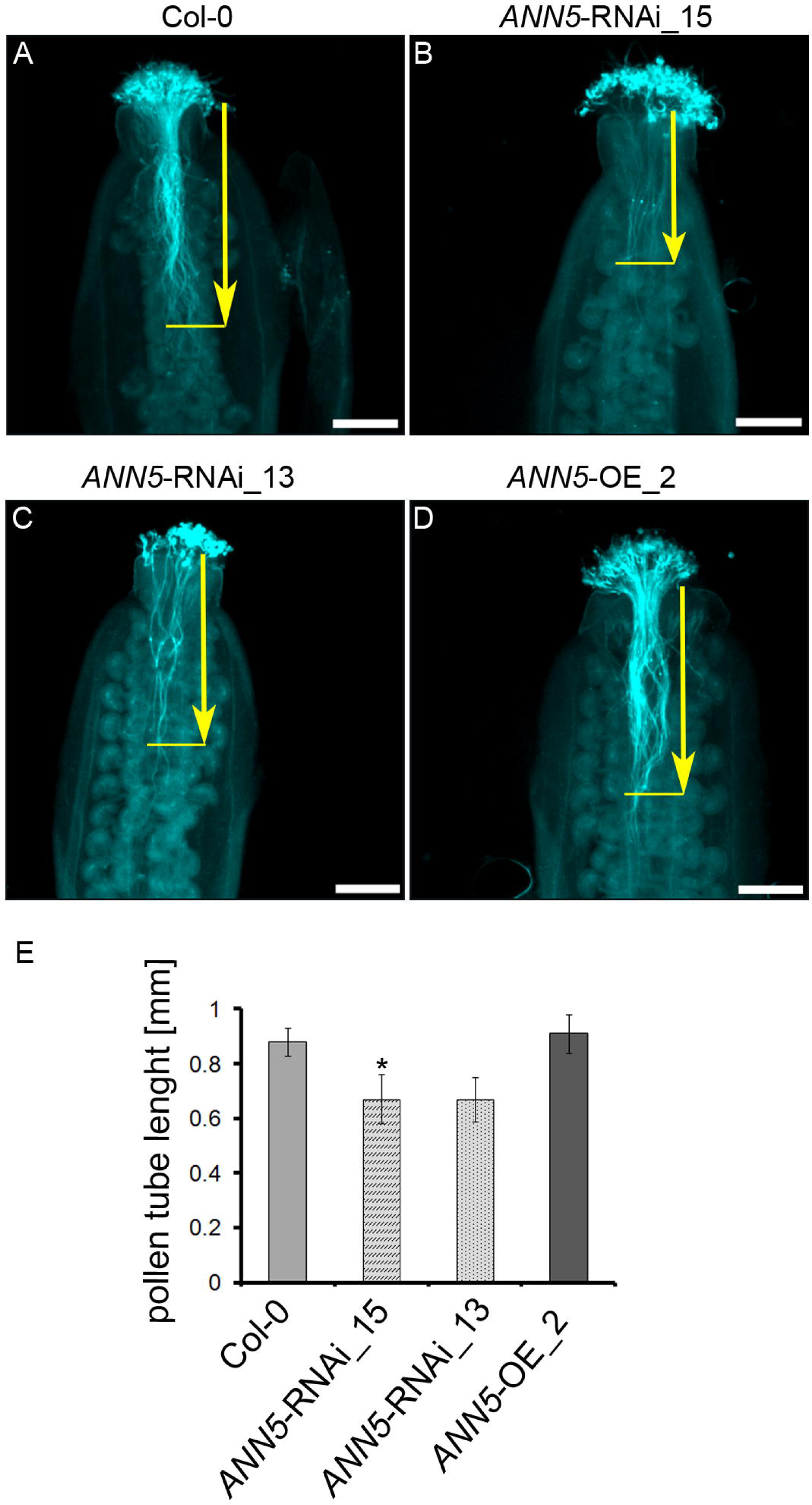
Pollen tube growth in pistils in *ANN5* RNAi-silenced lines. Pollen tubes were fixed and stained with Aniline Blue 6 h after hand-pollination of (A) wild-type Col-0, (B) *ANN5*-RNAi_15, (C) *ANN5*-RNAi_13, and (D) *ANN5-*OE_2 plants. Aniline blue staining of pollen tubes was performed as described by [51].Yellow arrows indicate pollen tube length measured from the top of style to the front of the longest pollen tube. (E) Average lengths of pollen tubes in pistils. n = 3 independent experiments. Asterisk indicates significant difference compared with the wild type (one-way ANOVA, Dunnett post hoc test, *p < 0.05; **p < 0.01; ***p < 0.001). See also Additional file 5. Scale bars = 200 μm.

### Total seed yield correlates with *ANN5* expression level

Although pollen viability was reduced, *ANN5* RNAi knockdown plants still produced sufficient amounts of viable pollen to successfully reproduce generatively. To quantify the final seed yield from lines with modified *ANN5* expression, 1000 seeds per genotype were collected and the individual seed areas were measured under a stereoscopic microscope. *ANN5* RNAi-silenced seeds were smaller, and *ANN5*-OE_2 seeds were larger, than wild-type Col-0 seeds (Fig. 4). To test whether silique position on the main bolt affected seed size, individual siliques were pooled into groups consisting of five successive siliques and the average seed size was calculated for each group. The average seed size decreased upwards towards the shoot in all the genotypes tested (Fig. 4C). Up to the 15th silique on the main bolt, seeds developed equally in wild-type Col-0 and *ANN5* RNAi knockdown plants. Above the 15th silique, average seed size was lower in *ANN5* RNAi lines than in the wild type. Average seed size decreased consecutively up the main bolt to the last examined silique, at the 40th node. Seeds collected from *ANN5*-OE_2 plants were consistently larger than wild-type seeds between the 11th and 40th nodes on the main bolt.

**Fig. 4.**
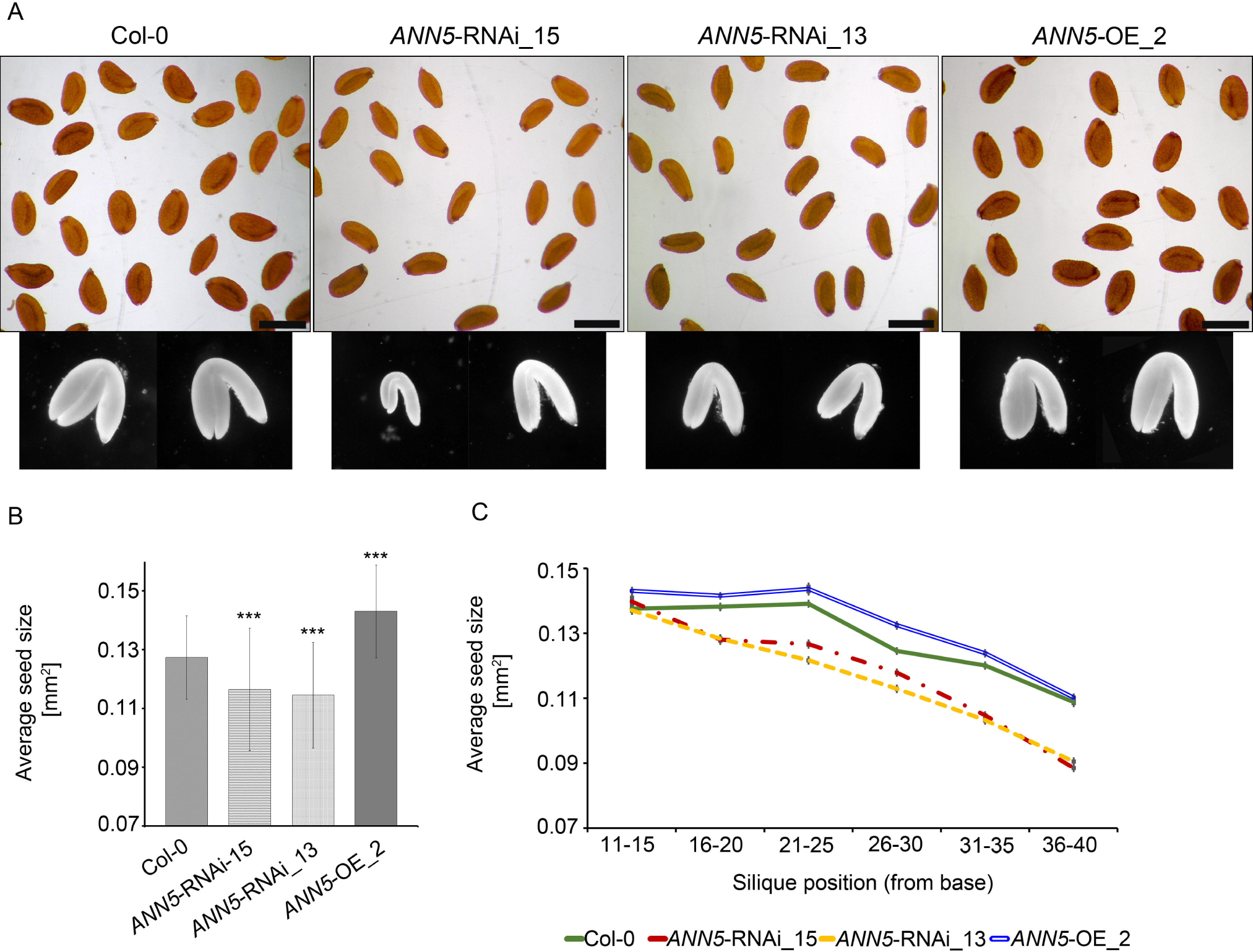
Impact of *ANN5* expression on seed yield. (A)Dry seeds isolated from siliques at positions 36–40 of the main bolt and embryos dissected from rehydrated seeds of wild-type Arabidopsis (Col-0), *ANN5*-RNAi_15, *ANN5*-RNAi_13, and *ANN5*-OE_2. Scale bars = 500 μm. (B)Average sizes of pooled seeds from a single biological replicate. n = 1000. Three independent experiments were performed with similar outcomes. Asterisks indicate significant differences compared with wild-type seeds (one-way ANOVA, Dunnettpost hoc test, *p < 0.05; **p < 0.01; ***p < 0.001). Bars represent SD. (C)Average sizes of seeds collected from siliques at specified positions on the main bolt, pooled from a single biological replicate. n = 120–150. Bars represent SD.

In Arabidopsis, embryos constitute most of the total volume of the mature seed, and the final size of dry seeds thus depends primarily on embryo size. Embryos dissected from *ANN5* RNAi-silenced seeds were smaller than those from wild-type seeds (Fig. 4A). Taken together, these results indicate that ANN5 affects flower and seed development during the reproductive phase of the Arabidopsis life cycle.

### Multi-compartment targeting of ANN5-GFP

Subcellular localization of ANN5 was analyzed to gain insights into the mechanisms underlying its functions. First, the online software tools PSORT and WoLF PSORT (www.genscript.com) were used to predict ANN5 subcellular localization. PSORT predicted localization to the nucleus, and WoLF PSORT predicted chloroplast localization. Additional software, Nuc-Plos, predicted ANN5 localization to the nucleolus.

Transient expression of *35S:ANN5-GFP* and *35S:GFP-ANN5* gene constructs in *Nicotiana benthamiana* leaves was used to examine subcellular localization of ANN5 *in vivo*. Confocal microscopy analysis revealed that ANN5-GFP was localized to the nucleus, nucleolus, and cytoplasm in all the epidermal cells examined (Fig. 5). In numerous cells, *ANN5-GFP* also accumulated in speckles inside the epidermal plastids, thus being fully consistent with the predictions.

**Fig. 5.**
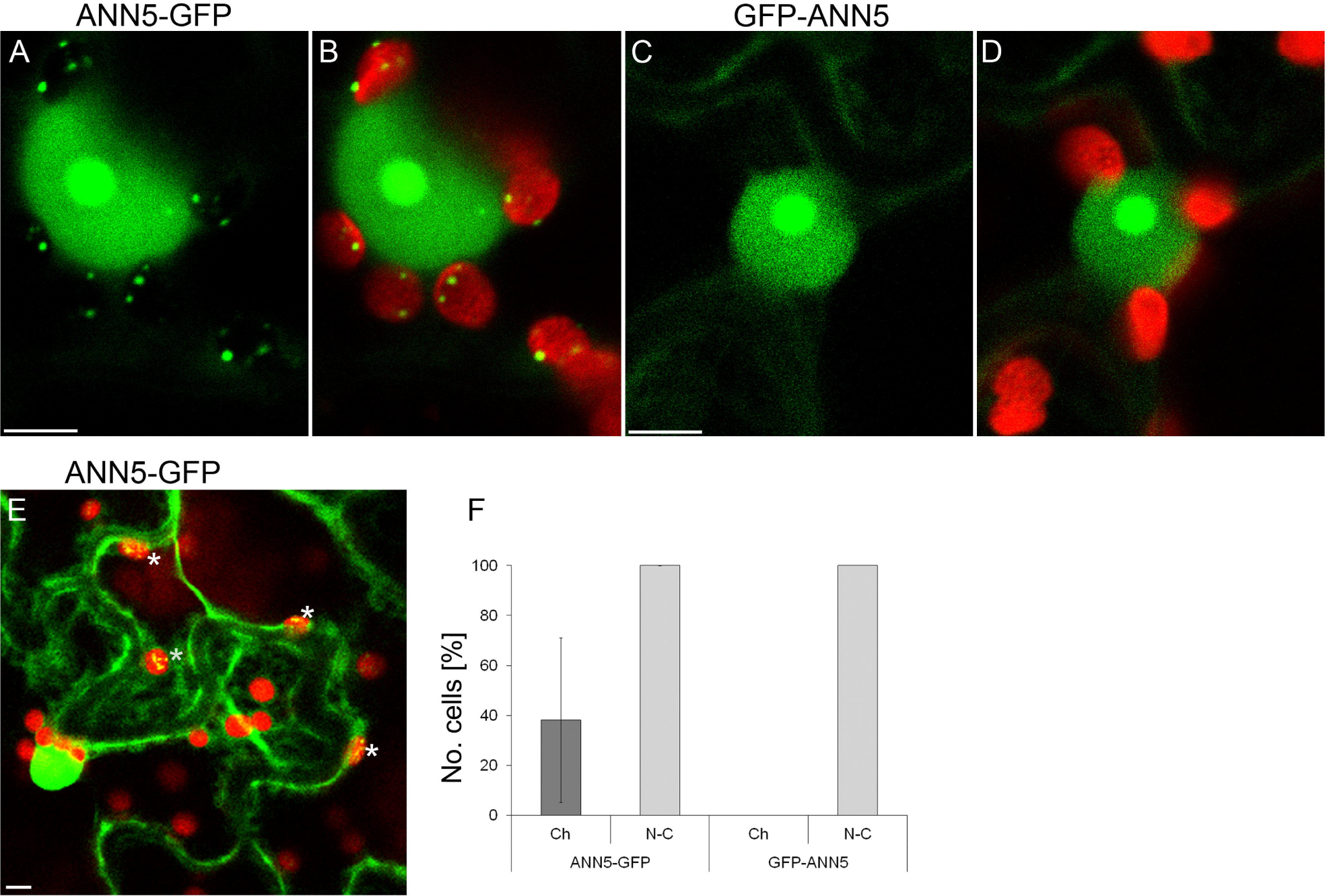
Subcellular localization of ANN5 in epidermal cells. Confocal optical sections of *N. benthamiana* leaf epidermal cells depicting localization of (A) C-terminus tagged ANN5 (35S:ANN5-GFP), (B) ANN5-GFP localization merged with chlorophyll autofluorescence, (C) N-terminus tagged ANN5 (35S:GFP-ANN5), and (D) GFP-ANN5 localization merged with chlorophyll autofluorescence. Scale bars = 10 μm. (E) Confocal optical section of two neighboring epidermal cells revealing different patterns of ANN5-GFP localization within plastids. White asterisks denote plastids containing ANN5-GFP. Scale bar = 10 μm. (F) Percentage of cells showing nucleo-cytoplasmic (N-C) or plastidial (Ch) localization of ANN5-GFP, and GFP-ANN5. The data were obtained in three independent experiments. Bars represent SD.

The number of cells in which ANN5-GFP localized to the plastids varied significantly between experiments. Notably, when ANN5 was found in a plastid within a cell, all the plastids of that cell contained ANN5 (Fig. 5h). N-terminal tagging with GFP resulted in the localization of ANN5 to the nucleus, nucleolus, and cytoplasm but eliminated plastid distribution (Fig. 5C and Fig. 5D). The punctate pattern of ANN5 distribution inside the plastids resembled the positioning of nucleoids. To test this, leaf samples expressing *35S:ANN5-GFP* were stained with DAPI: ANN5 speckles in plastids fully colocalized with DAPI-stained plastid DNA (Fig. 6).

**Fig. 6.**
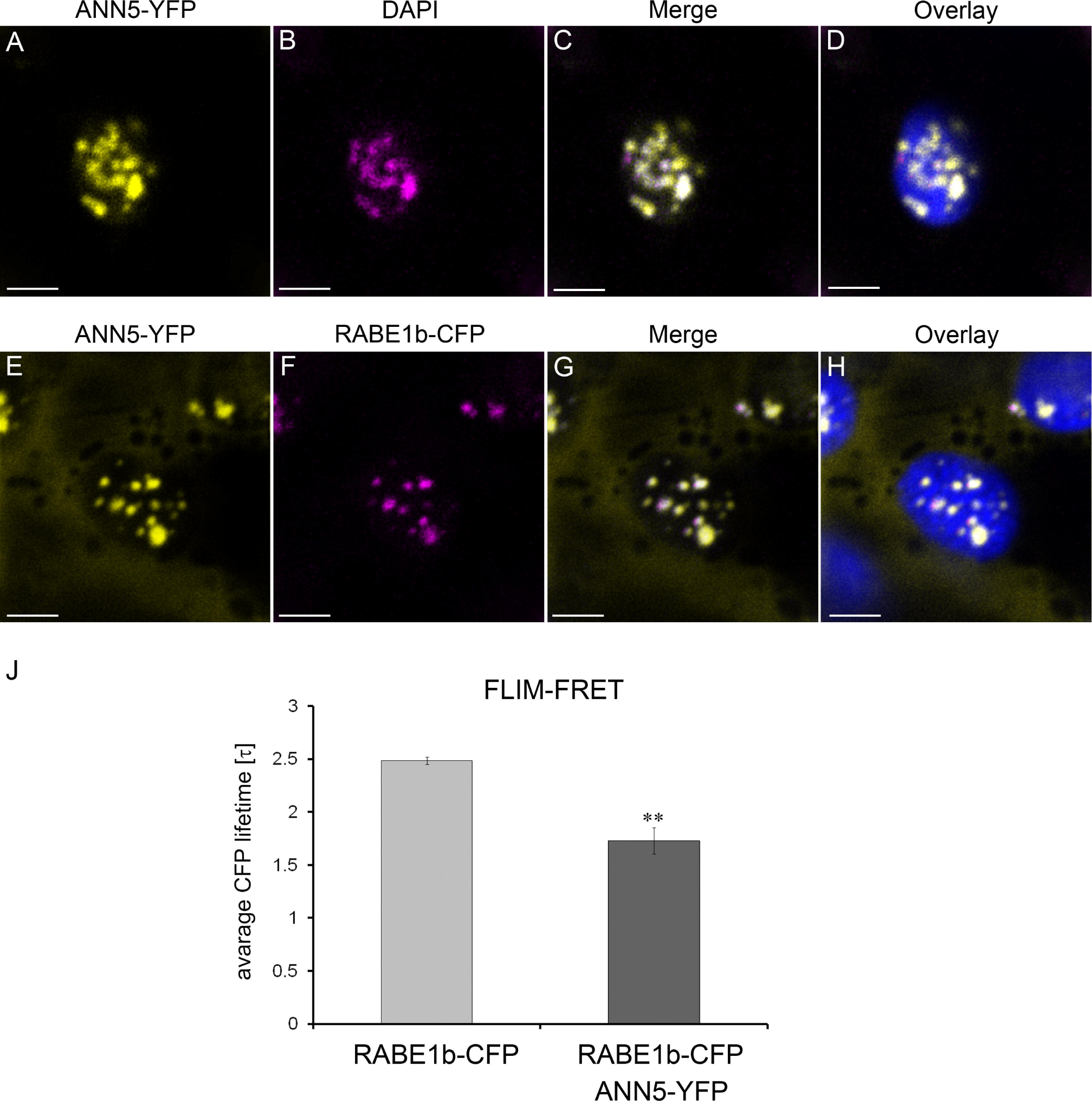
ANN5 interacts with RABE1b in plastidial nucleoids. Upper panel (A–B): Confocal optical section of *N. benthamiana* leaf epidermal plastid transiently expressing ANN5-YFP (A) and counterstained with DAPI (1 μg ml-1 for 15 min at room temperature) after fixation with 2% paraformaldehyde (24 h at 4°C) (B). Pseudocolored fluorescence of (A) ANN5-YFP (yellow), (B) DAPI (magenta), (C) merged channels of ANN5-YFP and DAPI, and (D) overlaid with chlorophyll autofluorescence (blue). Lower panel (E–H): Confocal optical section of *N. benthamiana* leaf epidermal plastid transiently co-expressing ANN5-YFP (E) and RABE1b-CFP (F). Pseudocolored fluorescence of (E) ANN5-YFP (yellow), (F) RABE1b-CFP (magenta), (G) merged channels of ANN5-YFP and RABE1b-CFP, and (H) overlaid with chlorophyll autofluorescence (blue). See also Additional file 7. Scale bar = 10 μm. (J) FLIM-FRET analysis of interactions between ANN5 and RABE1b in plastidial nucleoids. Average CFP lifetime was measured in the donor leaf samples of *N. benthamiana* expressing only RABE1b-CFP and in the presence of acceptor in samples co-expressing RABE1b-CFP and ANN5-YFP. n = 7 individual epidermal cells. Measurements were performed on a single plastid per cell, ** indicates statistically significant differences (Student’s t-test, p < 0.05).

### ANN5 interacts with RABE1b in plastid nucleoids

To identify ANN5 binding partners, 12-day-old Arabidopsis seedlings expressing *35S:ANN5-YFP* were used in co-immunoprecipitation experiments using a GFP-TRAP system followed by mass spectrometry. Identified proteins were compared between ANN5-YFP samples and control GFP samples, and proteins that non-specifically co-purified with GFP were excluded. Many of the identified proteins were predicted to be localized in plastids, suggesting that many of these associations might occur in plastidial nucleoids. To further investigate the specific interactions of ANN5 in plastids, potential binding partners were identified from proteins predicted to be localized in plastids (Additional file 6). Of these, RABE1b, which had the highest Mascot score and was a putative GTPase predicted to be plastid associated, was selected for further characterization.

Transient co-expression of *35S:ANN5-YFP* and *35S:RABE1b-CFP* in *N. benthamiana* was used to determine whether ANN5 and RABE1b localized to the same cellular compartment. When each was expressed alone, ANN5 localized to the nucleus, nucleolus, and plastid nucleoids (Fig. 5), and RABE1b-GFP was predominantly found within the plastid nucleoids and, to a lesser extent, in the cytoplasm (Additional file 7). When co-expressed, RABE1b-CFP and ANN5-YFP were detected within the same plastidial nucleoids (Fig. 6G). FLIM-FRET analysis was used to determine whether ANN5 and RABE1b interacted. In plastids, the average lifetime of the donor, RABE1b-CFP, decreased significantly in the presence of the putative acceptor ANN5-YFP (Fig. 6J). This confirmed physical interactions between ANN5 and RABE1b in the plastidial nucleoids.

### *ANN5* affects chlorophyll content in cotyledons of Arabidopsis seedlings

To check whether ANN5 affects plastid-related functions we analyzed greening of Arabidopsis seedlings with different *ANN5* expression levels. To this end, the seedlings were grown on MS medium in the absence or presence of sucrose, for ten days (Additional file 8, Fig. S7A). Whereas *ANN5* expression in wild-type seedlings was hardly detectable, the ectopic expression of *ANN5* resulted in abundant transcript levels (Additional file 8, Fig. S7C). Spectrophotometric analyses of chlorophyll a and b in the seedling revealed that the total chlorophyll content in *ANN5* OE lines was significantly lower than in wild-type seedlings on both types of media (Additional file 8, Fig. S7B). We next compared, the expression of selected genes related to the chlorophyll metabolism, by RT-qPCR analysis (Additional file 8, Fig. S7C and S7D). Both *ANN5* OE lines showed reduced expression of genes related to chlorophyll biosynthesis (*HEMA1, GUN4, GUN5*, C*HLI1*) and photosynthesis (*PsbA, LHCB1*) in comparison to the wild-type, whereas the expression of chlorophyll catabolic genes (*NYC1, NYE1, SAG29*) was higher but only in the presence of sucrose. These data show that *ANN5* overexpression affects chlorophyll accumulation in Arabidopsis seedlings.

## Discussion

### ANN5 plays an essential role during reproductive development of Arabidopsis

Annexins are implicated in a variety of cellular processes associated with membrane trafficking and calcium signaling [9-11, 25]. Annexins are mainly distributed within the cytoplasm and can reversibly interact with membranes in response to fluctuations in cellular calcium levels. When bound to calcium, hydrophobic residues are accessible on the surface of annexin, enabling interaction with phospholipids at the membrane interface [26]. ANN5 was also shown to possess the ability to bind lipids in a calcium-dependent manner [13]. Gradual increases in calcium concentration up to 200 μM enhanced ANN5 binding to liposomes *in vitro* [13]. Overexpression of *ANN5* in pollen tubes also conferred resistance to BFA, an inhibitor of the vesicular transport. Taken together, this research suggests that the biological activity of ANN5 might be related to membrane trafficking in a calcium-dependent manner.

The results from this study provide new insights into the function of ANN5 during Arabidopsis development. Large quantitative differences in *ANN5* transcript accumulation were observed between organs of wild-type Arabidopsis (Fig. 1), with the highest mRNA levels found in mature pollen. These results were consistent with a previous study showing that RNAi-mediated suppression of *ANN5* affected pollen development and led to reduced pollen viability [8]. Viable pollen grains from our RNAi knockdown lines were smaller in size and their growth in the pistil was hampered when compared with wild-type pollen grains (Fig. 2 and Fig. 3). In addition to its role in pollen grain development, through phenotypic studies, we showed that ANN5 was also involved in both embryo development and the transition from vegetative to generative growth (Table 1, Fig. 4). Suppression of *ANN5* resulted in extended vegetative development and reduced embryo size, whereas constitutive overexpression of *ANN5* positively influenced both pollen and embryo sizes. We thus conclude that ANN5 promotes cell growth, predominantly during the reproductive development of Arabidopsis.

### An insight into the role of ANN5 in plastid function

ANN5 displayed an unusual pattern of subcellular localization compared with the predominantly cytosolic localization observed for other plant annexins [11]. ANN5 occupied two DNA-containing cellular compartments (nucleus and plastid) and associated with prominent sub-organellar structures (nucleolus and plastidial nucleoids) (Fig. 5 and Fig. 6). The plastidial localization of ANN5 in a subset of cells suggested that ANN5 was mobile and might traffic to the plastids. N-terminal tagging of ANN5 with GFP inhibited its targeting to plastids while its nuclear distribution remained unaffected (Fig. 5). This confirmed that the N-terminal domain was essential for ANN5 import to the plastids. Moreover, mass spectrometry analysis of the C-terminal GFP fusion of ANN5 detected the peptide derived from the N-terminal region, suggesting that this signal was not cleavable. However, a scenario in which nuclear import of ANN5 does not require processing but import into the plastids requires cleavage of the N-terminal signal peptide cannot be excluded. This scenario would imply that transport of ANN5 from the nucleus to plastids is unidirectional or, alternatively, that the N-terminal sequence is protected from cleavage in the plastids, thus allowing shuttling of ANN5 between compartments.

Plastids are plant-specific organelles that possess their own genome and complete gene expression system [27]. Each type of plastid, except gerontoplasts, contains multiple copies of plastidial DNA arranged into compact structures termed nucleoids. Plastid nucleoids contain RNA and a multitude of proteins involved in the maintenance of nucleoid functions such as transcription, replication, RNA processing, and ribosome assembly [28, 29]. However, the majority of the proteins required for proper plastid function are encoded by the nuclear genome. Regulation of plastid functions is therefore continuously coordinated with the activity of the nuclear genome. An increasing body of evidence suggests that many nuclear proteins are also targeted to the plastids. The mechanism of dual targeting for many proteins is unclear. However, previous studies suggested that dual targeting might be either simultaneous or sequential [30]. Proteins that were initially targeted to the plastids and subsequently relocated to the nucleus might have a role in retrograde signaling. This mechanism of translocation was recently confirmed for HEMERA/pTAC12, which was targeted first to plastids and, after cleavage of its transit peptide, was relocated to the nucleus [31]. Our results suggest that ANN5 is localized primarily to the nucleus and then relocates to plastids. We hypothesize that ANN5 translocates from the nucleus directly to the plastidial nucleoid and then modifies plastid functions. Consistent with this model ANN5 negatively affected chlorophyll content and expression of the genes related to chlorophyll metabolism (Additional file 8). Principal component analysis (PCA) performed on expression levels of these genes, showed visible discrimination between groups corresponding to Col-0 and *ANN5*-overexpressing lines, suggesting a global influence of ANN5 presence on chlorophyll metabolism. Overexpression of *ANN5* resulted in the reduced expression of genes involved in chlorophyll metabolism e.g. *HEMA1, GUN4, GUN5*, C*HLI1, PsbA, LHCB1* and consequently in lower chlorophyll content. The fact that expression of the genes examined is sensitive to plastid signals [32-36] suggests that ANN5 is involved in communication between plastid and the nucleus. Interestingly, the addition of sucrose to the growth medium up-regulated genes associated with chlorophyll degradation in *ANN5*-overexpressing lines (*NYC1, NYE1, SAG29*) that implicates ANN5 in sucrose signaling pathway. Further work is needed to indentify the specific signals that drive ANN5-dependent reprogramming of plastid function. Recent studies revealed that retrograde regulation of the nuclear gene expression involved calcium signaling [37]. Calcium ions were released from the plastids to the cytosol in response to specific stimuli [38]. Cytosolic calcium transients were mediated by a plastid-localized calcium-sensing receptor, CAS. This process activated a MAP (mitogen-activated protein) kinase cascade, which in turn regulated activity of transcription factor ABI4 in the nucleus. The pattern of ANN5 subcellular distribution together with its calcium-dependent lipid-binding capacity might reflect its role in the crosstalk between the nucleus and plastids or in intraorganellar calcium signaling. Notably, previous research showed that intracellular redistribution of annexins in response to particular environmental stimuli was induced by calcium transients in the cytosol [39].

In summary, we hypothesize that ANN5 acts as a specific calcium signature decoder and orchestrates plastidial and nuclear genome activities in response to developmental and environmental cues. Disturbed bilateral communication between the nucleus and plastids might explain the retardation of reproductive development in *ANN5* RNAi-silenced plants. However, our hypothesis that the intracellular redistribution of ANN5 is calcium-dependent requires experimental verification.

During plastid differentiation, nucleoids undergo intensive remodeling and changes in their spatial arrangement but remain associated with the plastidial internal membrane [29]. Although poorly developed, the internal membrane system of Arabidopsis pollen grain plastids is thought to be photosynthetically active [2]. Since both pollen and embryo are sink organs that take up nutrients from other parts of the plant, their photosynthetic structures might be associated with processes other than conversion of light energy into sugars. Photosynthetic complexes in pollen grain plastids might act similarly to embryos and generate reactive oxygen species to regulate processes both inside plastids and, in response to the retrograde signaling, in the nucleus [40-42]. Recent studies suggested that plastidial nucleoids acted as a docking platform for the proteins involved in plastid metabolism that were regulated by redox changes in the photosynthetic apparatus [43]. One can thus speculate that ANN5, by combining membrane- and calcium ion-binding capacities, might act at the interface between the nucleoids and plastidial internal membranes. The majority of plastid/nucleus-targeted proteins were shown to be involved in plastid DNA/RNA metabolism or translation [44]. We therefore propose that ANN5 association with membrane-bound nucleoids may be required for transmission of signals from the photosynthetic apparatus to the transcription/translation machinery of the plastid.

*ANN5* expression correlated with post-meiotic development of microspores, which was accompanied by substantial reorganization of the plastid function. *ANN5* promoter activity was observed in the bicellular microspore [8], whereas *ANN5* mRNA levels were at their maximum in the tricellular microspore and remained high in mature pollen [7]. At the initial stage of pollen grain development, plastids intensively accumulate the starch that is deposited until the bicellular stage of microspore development [45]. From this stage until pollen grain maturity, deposited starch grains are almost completely hydrolyzed. Previous studies reported that plastids generated energy via glycolysis to support pollen maturation and pollen tube growth [6]. These findings together with our observation that ANN5 localized to plastids and affected the expression of the nuclear genes encoding plastid proteins raises the possibility that ANN5 may be involved in plastid reorganization at later stages of pollen development. Starch grains accumulated in aborted pollen grains of the *ANN5* RNAi_15 line. This suggested that abortion of pollen grains occurred at the bicellular stage, which was consistent with previous studies [8]. We thus conclude that suppression of *ANN5* disables progression to the next developmental stage and finally leads to pollen abortion at the bicellular stage (Fig. 7). Although average pollen grain size was significantly reduced in *ANN5* RNAi lines, individual pollen grains developed without any obvious aberrations (Fig. 2), possibly because *ANN5* was not completely suppressed (Fig. 2B). The *ANN5* knockdown phenotype resembled the phenotypes of Arabidopsis mutant lines defective in genes related to plastid function, including plastid glycolysis, that affected pollen formation, pollen tube growth, and embryogenesis [6, 46, 47]. Suppression of *ANN5* likely leads to plastid malfunction and, in turn, may affect the energy status of the cell and consequently lead to reduced growth or collapse of cells.

**Fig. 7.**
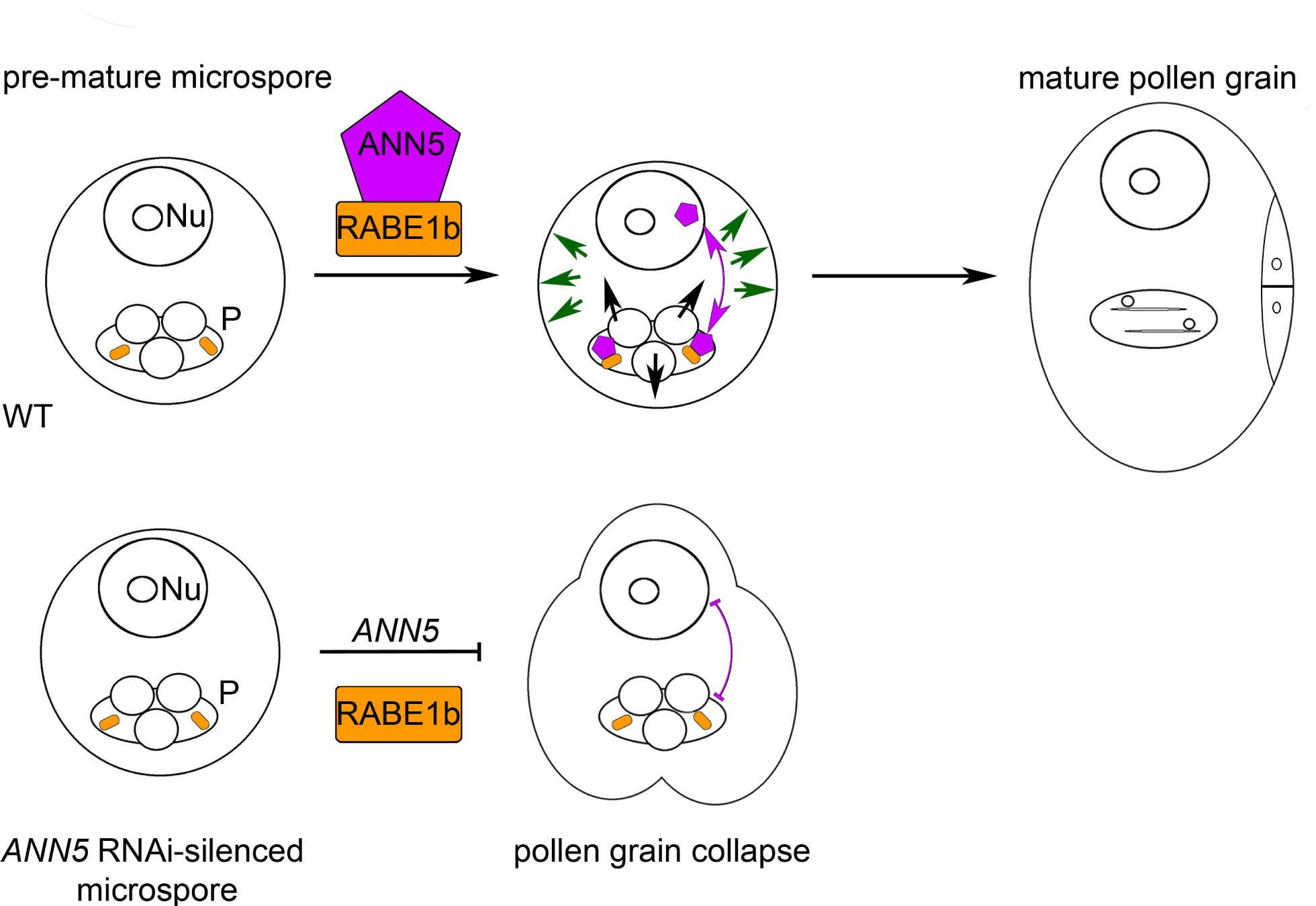
Hypothetical model for the role of ANN5 in pollen development. *ANN5* expression appears to be precisely temporally controlled during microgametophyte development in Arabidopsis. Onset of *ANN5* expression occurs in the bicellular pollen grain and remains expressed until maturation. Kinetics of *ANN5* expression correlate with reorganization of pollen plastid functions followed by gradual hydrolysis of deposited starch grains and progressive growth of the vegetative cell. ANN5 is dually located within the nucleus and plastidial nucleoids and may thus be involved in the crosstalk between nuclear and plastidial genomes (see also Additional file 8). Suppression of *ANN5* expression results in arrested plastid reorganization followed by pollen abortion. ANN5 interacted with RABE1b, a putative translational GTPase, within plastidial nucleoids. This model proposes that the physical interactions between ANN5 and RABE1b trigger a reprogramming of plastid function that is critical for proper pollen maturation. Disorder in cellular metabolism in genotypes with silenced expression of *ANN5* results in formation of smaller pollen grains and lower pollen viability. Nu: nucleus, P: plastid.

### Importance of the interaction between ANN5 and RABE1b for plastid functions

A large number of predicted plastid-targeted proteins were identified that co-purified with ANN5, including RABE1b, GAPA and GAPB subunits of glyceraldehyde 3-phosphate dehydrogenase (GAPDH), plastid chaperones, and ribosomal proteins (Additional file 6). Further characterization of RABE1b revealed physical interactions with ANN5 within plastidial nucleoids (Fig. 6J). Although the biological function of RABE1b is unknown, the protein contains a GTPase domain and is classified as a member of the Rab GTPase family, suggesting that it may be involved in intracellular trafficking [48]. Several proteins involved in the transport machinery were predicted to be plastid targeted, raising suggestions of vesicular transport within plastids [49].Given that both annexins and Rab GTPases are implicated in membrane trafficking, it is plausible that both ANN5 and RABE1b are required to maintain the organization and function of plastidial nucleoids attached to the plastid internal membranes.

RABE1b also exhibits sequence similarities to translation elongation factor EFTu/EF1A (www.arabidopsis.org) therefore it is likely that belongs to the translational GTPases [50]. In our hypothetical model for ANN5 function, we propose that ANN5 interaction with RABE1b occurs in the plastid nucleoids in the bicelluar microspore (Fig. 7). We hypothesize that cooperative action of ANN5 and RABE1b may drive the reprogramming of plastid function in maturing pollen grain. Further studies are required to elucidate the interplay between ANN5 and RABE1b in plastidial nucleoids and to determine whether their functions are associated with DNA/RNA metabolism or protein synthesis.

## Conclusions

Collectively, through this work, we showed that ANN5 was required for basal developmental processes during the transition from vegetative to generative growth and for pollen and embryo development. ANN5 likely accomplishes these activities through its membrane trafficking function in the nucleus and plastidial nucleoids. Our future work will focus on how the interaction between ANN5 and RABE1b could influence plastid functions, particularly during pollen grain development.

## Abbreviations

FLIM-FRET: Fluorescence Lifetime Imaging Microscopy-Förster Resonance Energy Transfer
ld: long day
OE: overexpression
PCA: principal component analysis
RNAi: RNA interference
sd: short day
SD: standard deviation
VC: vegetative cell

## Declarations

### Ethics approval and consent to participate

Not applicable

### Consent for publication

Not applicable.

### Availability of data and materials

All data generated or analyzed during this study are included in this published article and its supplementary information files. Arabidopsis accessions: Col-0, An-1, Bay-0, C24, Ler-1, Mr-0, Oy-0 and Wa-1 were obtained from NASC (http://arabidopsis.info/). Agrikola binary plasmids (http://www.agrikola.org/) for targeted *ANN5* RNAi silencing were obtained from NASC (http://arabidopsis.info/). Arabidopsis lines generated in this study and materials integral to the findings presented in this article are available on request at the Institute of Biochemistry and Biophysics, Polish Academy of Science in Warsaw (Poland) in the laboratory of corresponding author Malgorzata Lichocka.

### Author Contribution

M.L. and J.H. conceived and directed the research. M.L., W.R., M.K. and J.H. designed the experiments. M.L., W.R., K.M., I.B.F., A.Ch., M.S., E.S. performed research and analyzed data. M.L., M.A.S., M.K. and J.H wrote the paper. All authors have read and approved the final version of the manuscript.

## Acknowledgements

Imaging experiments were carried out using CePT infrastructure financed by the European Union’s European Regional Development Fund (Innovative Economy 2007-2013, Agreement POIG.02.02.00-14-024/08-00).

## Additional files

**Additional file 1.** Analysis of *ANN5* transcript abundance in flowers at anthesis collected from *ANN5* RNAi-silenced and overexpressing lines.

**Additional file 2.** Phenotypic characteristics of Arabidopsis with altered *ANN5* expression cultivated under a 12 h light regime.

**Additional file 3.** Impact of RNAi-mediated suppression of *ANN5* on pollen viability.

**Additional file 4.** Ultrastructure of bicellular microgametophytes isolated from Arabidopsis lines with altered *ANN5* expression.

**Additional file 5.** Growth of pollen tubes in pistils 24 h after hand-pollination.

**Additional file 6.** List of plastidial proteins co-purified with ANN5-YFP and identified by mass spectrometry.

**Additional file 7.** Subcellular localization of RABE1b-GFP in *N. benthamiana* leaf epidermal cells.

**Additional file 8.** Overexpression of *ANN5* influences chlorophyll content and alters expression of genes related to chlorophyll metabolism in Arabidopsis seedlings.

**Additional file 9.** Oligonucleotides used for RT-qPCR.

## References

1. Owen HA, Makaroff CA: Ultrastructure of microsporogenesis and microgametogenesis in Arabidopsis thaliana (L.) Heynh. ecotype Wassilewskija (Brassicaceae). Protoplasma 1995(185):7–21.

2. Kuang A, Musgrave ME: Dynamics of vegetative cytoplasm during generative cell formation and pollen maturation in Arabidopsis thaliana. Protoplasma 1996, 194:81–90.

3. Pacini E, Guarnieri M, Nepi M: Pollen carbohydrates and water content during development, presentation, and dispersal: a short review. Protoplasma 2006, 228(1-3):73–77.

4. Carrizo Garcia C, Nepi M, Pacini E: It is a matter of timing: asynchrony during pollen development and its consequences on pollen performance in angiosperms-a review. Protoplasma 2016.

5. Franchi G, Bellani L, Nepi M, Pacini E: Types of carbohydrate reserves in pollen: localization, systematic distribution and ecophysiological significance. Flora 1996, 191:143–159.

6. Selinski J, Scheibe R: Pollen tube growth: where does the energy come from? Plant Signal Behav 2014, 9(12):e977200.

7. Rutley N, Twell D: A decade of pollen transcriptomics.Plant Reprod 2015, 28(2):73–89.

8. Zhu J, Yuan S, Wei G, Qian D, Wu X, Jia H, Gui M, Liu W, An L, Xiang Y: Annexin5 is essential for pollen development in Arabidopsis. Mol Plant 2014a, 7(4):751–754.

9. Konopka-Postupolska D, Clark G: Annexins as Overlooked Regulators of Membrane Trafficking in Plant Cells. Int J Mol Sci 2017, 18(4).

10. Davies JM: Annexin-Mediated Calcium Signalling in Plants. Plants (Basel) 2014, 3(1):128-140.

11. Laohavisit A, Davies JM: Annexins. New Phytol 2011, 189(1):40–53.

12. Laohavisit A, Brown AT, Cicuta P, Davies JM: Annexins: Components of the calcium and reactive oxygen signaling network. Plant physiology 2010, 152(4):1824–1829.

13. Zhu J, Wu X, Yuan S, Qian D, Nan Q, An L, Xiang Y: Annexin5 plays a vital role in Arabidopsis pollen development via Ca2+-dependent membrane trafficking. PLoS One 2014 Jul 14;9(7):e102407 doi: 101371/journalpone0102407 eCollection 2014 2014b.

14. Herridge R, Day R, Baldwin S, Macknight R: Rapid analysis of seed size in Arabidopsis for mutant and QTL discovery. Plant Methods 2011, 7(1):3.

15. Gosh A, Pareek A, Singla-Pareek SL: Leaf Disc Stress Tolerance Assay for Tobacco. Bio-protocol 2015, 5(7).

16. Lee LY, Fang MJ, Kuang LY, Gelvin SB: Vectors for multi-color bimolecular fluorescence complementation to investigate protein-protein interactions in living plant cells. Plant Methods 2008, 4:24.

17. Nakagawa T, Suzuki T, Murata S, Nakamura S, Hino T, Maeo K, Tabata R, Kawai T, Tanaka K, Niwa Y et al: Improved Gateway binary vectors: high-performance vectors for creation of fusion constructs in transgenic analysis of plants. Biosci Biotechnol Biochem 2007, 71(8):2095–2100.

18. Hilson P, Allemeersch J, Altmann T, Aubourg S, Avon A, Beynon J, Bhalerao RP, Bitton F, Caboche M, Cannoot B et al: Versatile gene-specific sequence tags for Arabidopsis functional genomics: transcript profiling and reverse genetics applications. Genome Res 2004, 14(10B):2176–2189.

19. Clough SJ, Bent AF: Floral dip: a simplified method for Agrobacterium-mediated transformation of Arabidopsis thaliana. Plant J 1998, 16(6):735–743.

20. Lu Y: RNA Isolation from Arabidopsis Pollen Grains. Bio-protocol http://wwwbio-protocolorg/e67 2011, Bio101: pe67.

21. Giska F, Lichocka M, Piechocki M, Dadlez M, Schmelzer E, Hennig J, Krzymowska M: Phosphorylation of HopQ1, a Type III Effector from Pseudomonas syringae, creates a binding site for host 14-3-3 proteins. Plant physiology 2013, 161(4):2049–2061.

22. Golinowski W, Grundler FMW, Sobczak M: Changes in the structure of Arabidopsis thaliana during female development of the plant-parasitic nematode Heterodera schachtii. Protoplasma 1996, 194(1-2):103–116.

23. Kleemann J, Rincon-Rivera LJ, Takahara H, Neumann U, Ver Loren van Themaat E, van der Does HC, Hacquard S, Stuber K, Will I, Schmalenbach W et al: Sequential delivery of host-induced virulence effectors by appressoria and intracellular hyphae of the phytopathogen Colletotrichum higginsianum. PLoS Pathog 2012, 8(4):e1002643.

24. Clark GB, Sessions A, Eastburn DJ, Roux SJ: Differential expression of members of the annexin multigene family in Arabidopsis. Plant physiology 2001, 126(3):1072–1084.

25. Konopka-Postupolska D, Clark G, Hofmann A: Structure, function and membrane interactions of plant annexins: An update. Plant Science 2011, 181(3):230–241.

26. Lizarbe MA, Barrasa JI, Olmo N, Gavilanes F, Turnay J: Annexin-phospholipid interactions. Functional implications. Int J Mol Sci 2013, 14(2):2652–2683.

27. Sato S, Nakamura Y, Kaneko T, Asamizu E, Tabata S: Complete structure of the chloroplast genome of Arabidopsis thaliana. DNA Res 1999, 6(5):283–290.

28. Majeran W, Friso G, Asakura Y, Qu X, Huang M, Ponnala L, Watkins KP, Barkan A, van Wijk KJ: Nucleoid-enriched proteomes in developing plastids and chloroplasts from maize leaves: a new conceptual framework for nucleoid functions. Plant Physiol 2012, 158(1):156–189.

29. Powikrowska M, Oetke S, Jensen PE, Krupinska K: Dynamic composition, shaping and organization of plastid nucleoids. Front Plant Sci 2014, 5:424.

30. Krause K, Krupinska K: Nuclear regulators with a second home in organelles. Trends Plant Sci 2009, 14(4):194–199.

31. Nevarez PA, Qiu Y, Inoue H, Yoo CY, Benfey PN, Schnell DJ, Chen M: Mechanism of Dual Targeting of the Phytochrome Signaling Component HEMERA/pTAC12 to Plastids and the Nucleus. Plant Physiol 2017, 173(4):1953–1966.

32. McCormac AC, Terry MJ: Light-signalling pathways leading to the co-ordinated expression of HEMA1 and Lhcb during chloroplast development in Arabidopsis thaliana. Plant J 2002, 32(4):549–559.

33. McCormac AC, Terry MJ: The nuclear genes Lhcb and HEMA1 are differentially sensitive to plastid signals and suggest distinct roles for the GUN1 and GUN5 plastid-signalling pathways during de-etiolation. Plant J 2004, 40(5):672–685.

34. Ikegami A, Yoshimura N, Motohashi K, Takahashi S, Romano PG, Hisabori T, Takamiya K, Masuda T: The CHLI1 subunit of Arabidopsis thaliana magnesium chelatase is a target protein of the chloroplast thioredoxin. J Biol Chem 2007, 282(27):19282–19291.

35. Brzezowski P, Sharifi MN, Dent RM, Morhard MK, Niyogi KK, Grimm B: Mg chelatase in chlorophyll synthesis and retrograde signaling in Chlamydomonas reinhardtii: CHLI2 cannot substitute for CHLI1. J Exp Bot 2016, 67(13):3925–3938.

36. Mochizuki N, Tanaka R, Tanaka A, Masuda T, Nagatani A: The steady-state level of Mg-protoporphyrin IX is not a determinant of plastid-to-nucleus signaling in Arabidopsis. Proc Natl Acad Sci U S A 2008, 105(39):15184–15189.

37. Guo H, Feng P, Chi W, Sun X, Xu X, Li Y, Ren D, Lu C, David Rochaix J, Leister D et al: Plastid-nucleus communication involves calcium-modulated MAPK signalling. Nat Commun 2016, 7:12173.

38. Stael S, Wurzinger B, Mair A, Mehlmer N, Vothknecht UC, Teige M: Plant organellar calcium signalling: an emerging field. J Exp Bot 2012, 63(4):1525–1542.

39. Clark GB, Morgan RO, Fernandez MP, Roux SJ: Evolutionary adaptation of plant annexins has diversified their molecular structures, interactions and functional roles. New Phytol 2012, 196(3):695–712.

40. Allorent G,Osorio S, Vu JL, Falconet D, Jouhet J, Kuntz M, Fernie AR, Lerbs-Mache S, Macherel D, Courtois F et al: Adjustments of embryonic photosynthetic activity modulate seed fitness in Arabidopsis thaliana. New Phytol 2015, 205(2):707–719.

41. Kim C, Lee KP, Baruah A, Nater M, Gobel C, Feussner I, Apel K: (1)O2-mediated retrograde signaling during late embryogenesis predetermines plastid differentiation in seedlings by recruiting abscisic acid. Proc Natl Acad Sci U S A 2009, 106(24):9920–9924.

42. Yoshida K, Hisabori T: Two distinct redox cascades cooperatively regulate chloroplast functions and sustain plant viability. Proc Natl Acad Sci U S A 2016, 113(27):E3967–3976.

43. Melonek J, Oetke S, Krupinska K: Multifunctionality of plastid nucleoids as revealed by proteome analyses. Biochim Biophys Acta 2016, 1864(8):1016–1038.

44. Krause K, Oetke S, Krupinska K: Dual targeting and retrograde translocation: regulators of plant nuclear gene expression can be sequestered by plastids. Int J Mol Sci 2012, 13(9):11085–11101.

45. Quilichini TD, Douglas CJ, Samuels AL: New views of tapetum ultrastructure and pollen exine development in Arabidopsis thaliana. Ann Bot 2014, 114(6):1189–1201.

46. Niewiadomski P, Knappe S, Geimer S, Fischer K, Schulz B, Unte US, Rosso MG, Ache P, Flugge UI, Schneider A: The Arabidopsis plastidic glucose 6-phosphate/phosphate translocator GPT1 is essential for pollen maturation and embryo sac development. Plant Cell 2005, 17(3):760–775.

47. Datta R, Chamusco KC, Chourey PS: Starch biosynthesis during pollen maturation is associated with altered patterns of gene expression in maize. Plant Physiol 2002, 130(4):1645–1656.

48. Vernoud V, Horton AC, Yang Z, Nielsen E: Analysis of the small GTPase gene superfamily of Arabidopsis. Plant Physiol 2003, 131(3):1191–1208.

49. Paul P, Simm S, Mirus O, Scharf KD, Fragkostefanakis S, Schleiff E: The complexity of vesicle transport factors in plants examined by orthology search. PLoS One 2014, 9(5):e97745.

50. Maracci C, Rodnina MV: Review: Translational GTPases. Biopolymers 2016, 105(8):463–475.

51. Mori T, Kuroiwa H, Higashiyama T, Kuroiwa T: GENERATIVE CELL SPECIFIC 1 is essential for angiosperm fertilization. Nat Cell Biol 2006, 8(1):64–71.

